# *LubriShield^TM^* - a unique permanent coating for indwelling urinary catheters that impedes surface-associated uropathogens from forming biofilm

**DOI:** 10.1101/2025.01.24.634505

**Authors:** Ana I. Romero, Serhiy Surkov, Per Wirsén, Graeme Brookes, Linda Bergström, Jan Tejbrant, Elena Dhamo, Sandra Wilks, Catherine Bryant, Jan Andersson

## Abstract

Catheter-associated urinary tract infection (CAUTI) is one of the most common healthcare-associated infections and biofilm formation plays a key role in its pathogenesis. Indwelling medical devices introduce ideal pathways inside the body for invading pathogens and feature surfaces conducive to biofilm development. These devices are the root cause of severe clinical infections often recalcitrant to antimicrobials. When bacteria and fungi switch to biofilm mode of growth, they produce a matrix in the form of extracellular polymeric substances (EPS). This creates a unique environment for growing virulent colonizers and persisting cells while also forming a shielding barrier against immune system attacks, antimicrobial agents and mechanical removal by fluid shear forces.

To address this challenge, LubriShield ^TM^ - a novel permanent coating was invented, and evenly applied to both the internal and external surfaces of indwelling urinary Foley catheters. Without releasing active substances, it effectively prevented pathogens from producing biofilm. The coating was superhydrophilic and incorporated a proprietary anti-fouling ligand, which created a surface that significantly inhibited up to 99% of colonizing uropathogens from forming biofilm for the duration of use without any microbial killing (p< 0.001). The predominant uropathogens, including Gram-positive and Gram-negative as well as Candida albicans, were inhibited from forming biofilm on the LubriShield^TM^ coated surfaces for up to 14 days in artificial urine medium. After challenging the adhering bacteria in a glass bladder flow model, the coating still significantly reduced biofilm formation by 83% (p<0.001).

The growth mode and emergent properties of adhering bacteria on uncoated silicone catheter surfaces were compared with those on LubriShield^TM^ coated surfaces. RNA-seq analysis revealed that gene expression associated with microbial EPS formation was significantly downregulated on the coated surfaces. Additionally, microorganisms adhering to LubriShield^TM^ coated catheters were 46% more susceptible to antibiotics compared to those on uncoated silicone catheters (p<0.01).

## Introduction

Among urinary tract infections (UTIs) acquired in the care of patients, approximately 75% are associated with the use of urinary catheters ^1^. Catheter-associated urinary tract infection (CAUTI) is usually caused by biofilms formed by microorganisms originating from the subject’s own colonic and perineal microflora (*Escherichia coli, Klebsiella pneumoniae, Proteus mirabilis, Pseudomonas aeruginosa, Enterococci*), but can also come from the hands of healthcare personnel or the patient’s skin during catheter insertion and manipulation with the collection system (*Staphylococcus epidermidis* or S*taphylococcus aureus*). CAUTIs may lead to potentially serious consequences, including frequent febrile episodes, acute and chronic pyelonephritis, bacteraemia with urosepsis and septic shock, catheter obstruction and renal and/or bladder stone formation, and even bladder cancer with prolonged use ^2–4^.

Catheters provide an ideal surface on which bacteria can adhere and develop biofilms ^5,6^. The catheter-associated biofilm is a three-dimensional structure triggered by the microorganism’s surface sensing and initial adhesion, followed by colonization and the production of a slimy matrix composed of extracellular polymeric substances (EPS) ^7^. It represents a predominant form of microbial life with highly organized communities of microorganisms, including bacteria, fungi, and, at times, mixed-species populations. Within a biofilm, distinct population zones emerge as a result of environmental gradients. Persistent cells often reside near the base of the biofilm in anaerobic regions, where nitrate respiration dominates in the absence of oxygen. In contrast, metabolically active, oxygen-respiring cells occupy the biofilm’s outer layers, poised for dispersal to new surfaces ^8^. Only approximately 10% of the biofilm mass represents microbial cells, while most of it consists of hydrated EPS. Polysaccharides, proteins and extracellular DNA (eDNA) are the main components of the EPS, but it can also contain varying amounts of membrane vesicles, lipids, amyloids, cellulose and appendages, such as cell-wall-anchored proteins, pili, fimbriae, and flagella. These components not only provide structural stability to the biofilm but can also contribute to inflammatory conditions ^9^.

Catheter-associated biofilms provide uropathogens with many survival advantages, including resistance to the shear stress of the urine flow, resistance to phagocytosis, and resistance to antimicrobial agents ^10–12^. In addition, some common uropathogens, such as *Proteus species*, *P. aeruginosa*, and *K. pneumoniae*, produce an active urease that can hydrolyse urea in the urine to free ammonia ^13^. This results in a rapid increase in local pH leading to precipitation of minerals, such as hydroxyapatite and/or struvite. Encrustations of this kind are seen typically on the inner lumen of the catheter and can build up to block the catheter flow completely ^14^. Despite continued efforts to produce effective catheter materials and coatings that resist biofilm development, the problems of CAUTI and blockage prevail. Coatings that kill bacteria by release only have limited effect in the immediate surroundings and exhaust after a while. To date, bactericidal and bacteriostatic materials, such as silver-coated or antibiotic-impregnated coatings, have not been found to improve clinical outcomes significantly over long-term use ^10,15,16^. Other attempts with antibacterial topological coatings that kill bacteria upon adhesion often get compromised by the accumulation of dead cells and associated debris on the surfaces ^17^. These coatings do not prevent biofilm formation, but promote antimicrobial resistance, generate endotoxin release from dying bacteria and are potentially toxic to the human body. Thus, there is an unmet clinical need for effective long-lasting, non-release protection against biofilm formation on indwelling catheters. By understanding how microorganisms sense a surface and what signals make them switch into a virulent lifestyle and biofilm mode of growth, more efficient coatings can be developed to protect against catheter-induced infections.

Here we describe a novel approach for a permanent non-release superhydrophilic surface coating. The LubriShield^TM^ coating consists of a covalently bound polyacrylic-acid hydrogel coupled with a proprietary anti-fouling ligand. The coating has demonstrated an impressive effect in both static and dynamic culture conditions inhibiting microbial biofilm formation along the inside and outside of the catheter. The LubriShield^TM^ coated CytaCoat Foley catheter is biocompatible, nontoxic and produces minimal friction. When wet, the coating reduces friction of the catheter surface up to 28-fold as compared to uncoated silicone foley catheters. Coated catheters were exposed to uropathogens (*E. coli*, *K. pneumoniae*, *P. mirabilis*, *P. aeruginosa*, *Enterococcus faecalis*, *S. epidermidis*, *S. aureus*, and *Candida albicans*) for up to 14 days in artificial urine and evaluated for inhibition of biofilm formation by morphological, biochemical and genetic data.

## Results

### The LubriShield^TM^ coated surface prevented multiple uropathogenic strains from forming biofilm

The ability of different strains of the most common uropathogens to form biofilm in artificial urine media (AUM) on urinary catheters was tested under static condition at 37° after 5 and up to 14 days. Coated and uncoated silicone urinary catheters were aseptically cut into 2-cm lengths and individual pieces were inoculated with 10^9^ CFU/mL of the different uropathogens in AUM. Culture media was replaced every day to provide fresh nutrients and to discard unattached bacteria. After 7 days, all uropathogens had formed thick biofilms on the control silicone catheter surfaces. Representative fluorescence microscopy images show that the LubriShield^TM^ coated catheters prevented adhering *K. pneumoniae*, *E. faecium*, and *C. albicans* producing detectable amounts of polysaccharides on the surface determined by staining with Alexa 549-coupled fluorescent concanavalin A (fig. 1A). Further staining with a non-specific protein fluorochrome, SYPRO Ruby, revealed that in contrast to the uncoated silicone catheter surface, no protein components of the EPS were found on the LubriShield^TM^ coated surface regardless of which uropathogenic microbe that was incubated in the AUM (fig. 1B).

**Figure 1.**
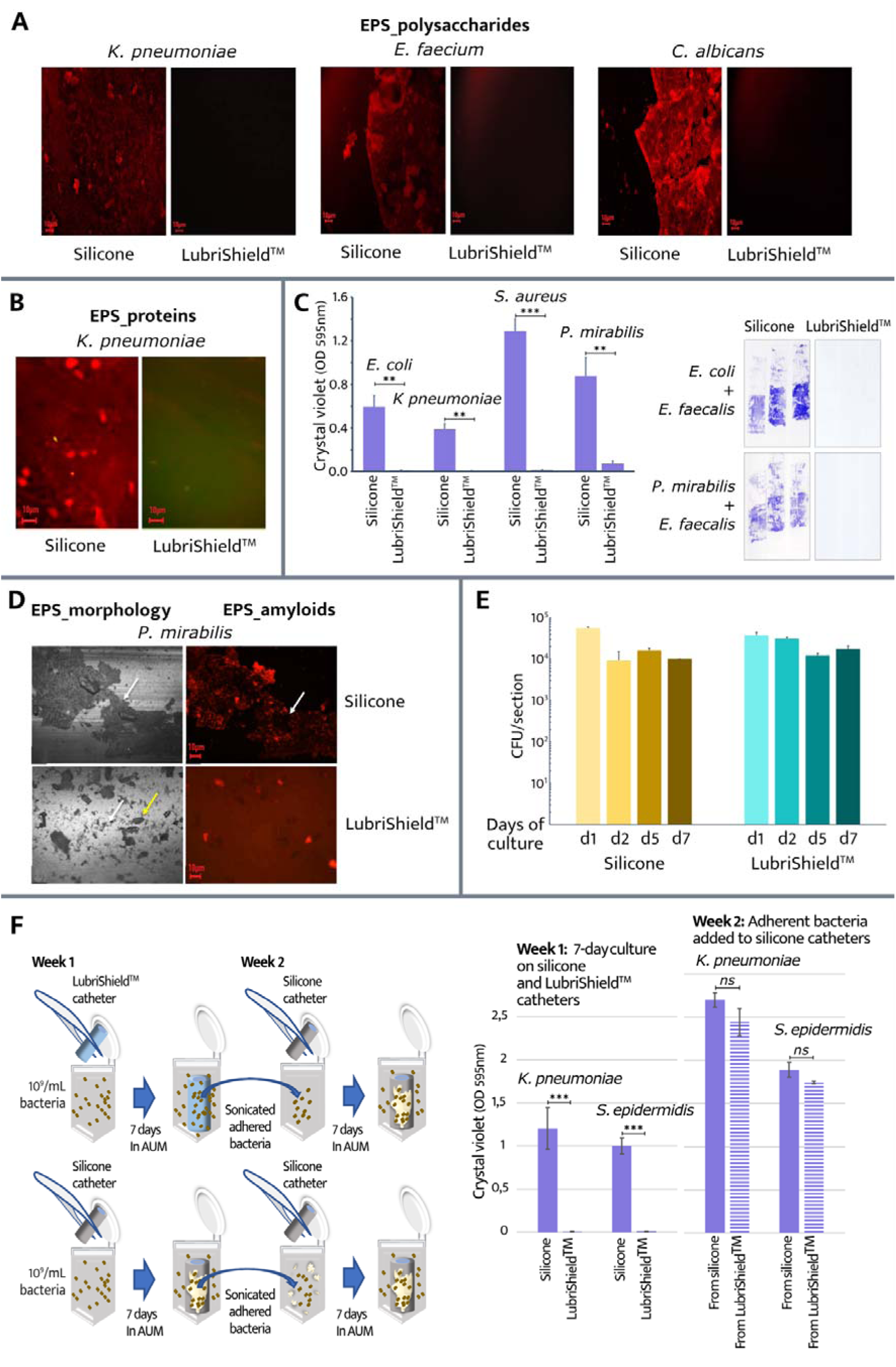
Assessment of long-term biofilm cultures in artificial urine of different uropathogenic strains on uncoated and LubriShield^TM^ coated silicone catheters. **A.** 7-day cultures in artificial urine under static conditions show significantly reduced production of polysaccharides by K. pneumoniae, E. faecalis or C. albicans on LubriShield^TM^ catheter surfaces. Microscopic images of concanavalin A fluorescent staining (Alexa Fluor™ 594 conjugate) comparing control silicone-versus LubriShield^TM^ catheter surfaces (repeated three times). **B.** Microscopic images of SYPRO Ruby fluorescent staining (FilmTracer ^TM^ SYPRO® Ruby) of proteins by K. pneumoniae comparing control silicone-versus LubriShield^TM^ catheter surfaces (repeated three times). **C.** Quantification of polysaccharides and proteins in the EPS by crystal violet shows no biofilm after 14 days of static culture with different uropathogens (Student’s t test: n=3, ***=p<0.001, **=p<0.01). The right panel shows crystal violet staining on adhesive tape transferred from 14 day mixed-species culture on control silicone-versus LubriShield^TM^ catheter surfaces. **D.** Images of parts of catheter surfaces after 5 days of incubation in AUM showing EDIC/EF images in the left panels and EbbaBiolight 680 staining of EPS in the right panels. Magnification = X100, White arrows = biofilm, Yellow arrows = crystal formation. **E.** Colony forming units (CFU) per section of P. mirabilis bacteria over time incubated in AUM comparing silicone catheter surface to LubriShield^TM^ surface (n=3). **F.** Schematic representation of the conducted experiment assessing biofilm-forming abilities of bacteria on LubriShield^TM^ surface (left). Crystal violet counts showing biofilm formation ofthe P. mirabilis residing on LubriShield^TM^ catheter and on silicone catheter surface (week 1) or removed from the corresponding surface and re-cultured on uncoated silicone (week 2 (n=3, *** = P<0.001, ns= nonsignificant, Student’s t test).

Concentrated inoculations with single and mixed strains of *E. coli, K. pneumoniae, S. aureus, P. mirabilis and E. faecalis*(10^9^ CFU/mL) were cultured on catheter pieces for 14 days in AUM. Quantification of the adherent biomass, including cells and EPS, was assessed by the crystal violet staining method. The absorbance of the extracted staining showed that still after 14 days, there was a significantly reduced biofilm formation for all tested uropathogens on the LubriShield^TM^ catheter surface compared to the silicone surface (fig. 1C) (ranging from p<0.01 to p<0.001). Scotch tape captures of the crystal violet staining showed how the LubriShield^TM^ catheter surface was clean even after inoculation with common mixed-population biofilm species such as *E. coli* and *E. faecalis* or *P. mirabilis* and *E. faecalis* (fig. 1C)

Uncoated and coated catheters were incubated with exponentially grown liquid culture of the uropathogenic *P. mirabilis* in AUM. There were no significant differences in pH values between the cultures containing the uncoated silicone and LubriShield^TM^ coated catheters. In both cases considerable increase to pH 9.0-9.5 was seen within 24 hours, and then maintained through the duration of the experiment. Catheter samples were also examined using EDIC/EF microscopy. EDIC imaging uses long working distance, non-contact, high magnification objectives to allow non-destructive imaging of biofilm structures and their interactions with surface materials ^18^. The appearance of crystals, particularly struvite’s, were observed in all samples as expected with the observed increase in pH. The internal and external catheter surfaces of the uncoated control sample showed a thick biofilm covering big areas in a sheath-like structure (fig. 1D). In contrast, the LubriShield^TM^ coated catheters showed a patchier, mosaic-like pattern, indicating microcolonies of bacteria. Assessing the biofilm structure using the EbbaBiolight stain, which labels amyloid proteins and negatively charged polysaccharides with repeating units in the EPS, showed only signal on the uncoated silicone surface as well as on crystalline formations that cause adsorption of the stain (fig. 1D).

### The LubriShield^TM^ coating did not exert any microbial killing effect

Quantification of the attached bacteria by culture techniques did not show any significant differences between the uncoated and coated samples as seen in figure 1E. In all cases, the colony-forming ability was maintained during the seven-day culture. This confirmed that there was no bactericidal or bacteriostatic effect of the LubriShield^TM^ coating (fig. 1E).

To analyse if the LubriShield^TM^ coating had a permanent biofilm-preventive effect on bacteria, uncoated and coated catheter pieces were inoculated with 10^9^ CFU/mL of either *K. pneumoniae* or *S. epidermidis* for 7 days in AUM with daily medium replacement. The catheter-adhering bacteria were sonicated off, counted and transferred to new tubes with uncoated catheter pieces and cultured again for seven days in AUM. Biofilm formation was assessed by staining all catheter surfaces with crystal violet and the extracted staining was then read in a spectrophotometer. The results showed that although biofilm formation by adhering bacteria was significantly inhibited on the LubriShield^TM^ coated catheter surfaces, the same bacteria were able to form biofilm on uncoated catheters equal to the biofilm-forming bacteria coming from the uncoated surfaces (fig. 1F).

### A strong anti-biofilm effect occurred even under dynamic conditions

We challenged the ability of uropathogens to form catheter-associated biofilms during dynamic conditions in a flow model. Uncoated and coated catheters were inoculated with urease-positive *P. mirabilis* (10^9^ CFU/mL) and incubated at 37°C under a pulsing flow of AUM for 3 days. At the end of the test, the control silicone catheter lumen was completely blocked with crystal precipitation in contrast to the LubriShield^TM^ coated catheter lumen, where less crystals had formed (fig. 2A). Labelling of EPS on the same catheter surfaces with EbbaBiolight 680, showed that the biofilm formation covered almost all of the control silicone catheter surface contrary to the LubriShield^TM^ coated catheter surface. Significantly less biofilm could be spotted (71% reduction, p<0.005) predominantly associated with biomass present on the surface of crystals (fig. 2A).

**Figure 2.**
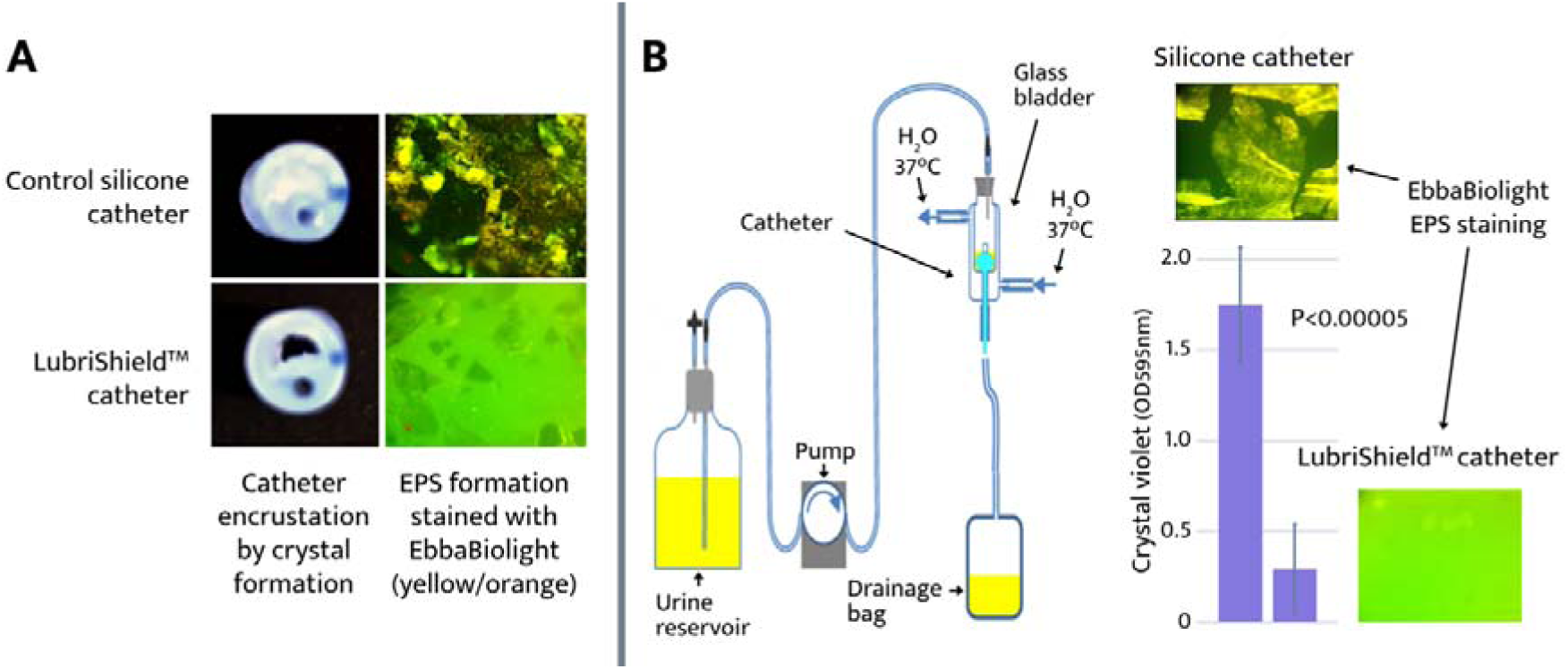
Biofilm assessment of the LubriShield^TM^ catheter in dynamic flow models. **A.** Crystal formation (left panels) and fluorescent labelling of biofilm (right panels, magnification X 40) on LubriShield^TM^ surface after P. mirabilis cultured in AUM for 3 days in a drip flow reactor. **B.** Schematic illustration of the in vitro glass bladder model system (left panel). Water was circulated through the outer chamber of a double walled vessel to maintain the glass bladder at 37°C. Catheters were inserted into the inner chamber and retention balloons inflated to hold the catheter in place. Catheters were connected to standard drainage bags below the level of the vessel to form a complete sterile closed drainage system. AUM was supplied to the glass bladder from the top at a constant flow rate controlled by a pump ^19^. Right panel shows crystal violet counts and fluorescent staining of the catheter sections from a glass bladder experiment after 5 days of P. aeruginosa culture in AUM (p<0.00005).

To further consider the hydrodynamic influence of the urine flow on the development of catheter-associated biofilms, we compared the whole length of the LubriShield^TM^ coated silicone catheters with uncoated catheters in an *in vitro* glass bladder model system (fig. 2B, left panel). Six bladders with three LubriShield^TM^ coated catheters, and three uncoated catheters were set up to study the biofilm formation of *P. aeruginosa*. A peristaltic pump provided AUM flow through the catheter at a rate of 1.5 mL/min and the temperature was held at 37°C. After five days, the models were stopped, and the catheters were analysed. Formation of biofilm on the luminal surfaces of the catheters was assessed by labelling EPS with EbbaBiolight 680 for microscopic determination and crystal violet staining for quantitative measurements. Results calculated from the mean crystal violet absorbance of three catheter replicates showed an 83% significant reduction (p<0.00005) in biomass on the LubriShield^TM^ coated catheter compared to the uncoated catheter (fig. 2B, right panel). Fluorescence microscopy imaging of the EPS formation confirmed the anti-biofilm effect of the LubriShield^TM^ coating (fig. 2B, right panel).

### Reduced biofilm-associated gene expression on the LubriShield ^TM^ coated catheter

Cells in biofilm mode manifest different gene expression pattern compared to the planktonic ones. *Pseudomonas aeruginosa* PAO1, a well-studied laboratory strain with a sequenced genome and often used as a model for biofilm-related gene analysis ^20,21^, was used to compare gene expression on mRNA level in bacteria growing for seven days on the LubriShield^TM^ coated surface in comparison to uncoated silicone catheter using RNA-seq analysis. Significant difference in the mRNA transcript was observed, with more than half of all the genes changing expression between the two groups (supplementary fig. S1).). We conducted analysis of the mRNA expression levels of key EPS-related operons. As figure 3A shows, the expression levels of several key genes in those operons were significantly higher on the un-coated surface compared to LubriShield^TM^. In particular the genes responsible for the production of alginate (*alg44* (PA3542), *algE* (PA3544), *algF* (PA3550), *algK* (PA3550), *algL* (PA3547); pel polysaccharide (*pelC* (PA3062), *pelG* (PA3058); psl polysaccharide (*pslM* (PA2243), *pslN* (PA2244); tight adherence proteins constituting type IVb pili (*tadA* (PA4302), *tadC* (PA4300), *tadD* (PA4299), *tadG* (PA4297)); fimbrial CU-pili adhesins (*cupA1* (PA2128)*, cupA3*(PA2130)*, cupA4*(PA2131)*, cupB1*(PA4086)*, cupC2*(PA0993)*, cupE4*(PA4651)*, cupE5*(PA4652)*, cupE6*(PA4653); and lectins, *lecA* (PA2570)) were found to be upregulated. The most striking difference was observed for the genes involved in the nitrate respiration, common among bacteria residing at the bottom of biofilms. As can be seen in figure 3b, 67% of the nitrate respiration genes had significantly higher mRNA expression levels on the uncoated silicone catheter surface compared to the LubriShield^TM^ coated one. In addition, genes involved in arginine fermentation, which is another sign of anaerobic lifestyle in *Pseudomonas,* as well as other hypoxia-related genes ^22^ had significantly higher levels on the uncoated silicone surface (supplementary table S1). This finding indicated that a significant proportion of the cells were in an anaerobic environment, a characteristic of complex biofilm structure, while no such structure was present on LubriShield^TM^ coated catheters.

**Figure 3.**
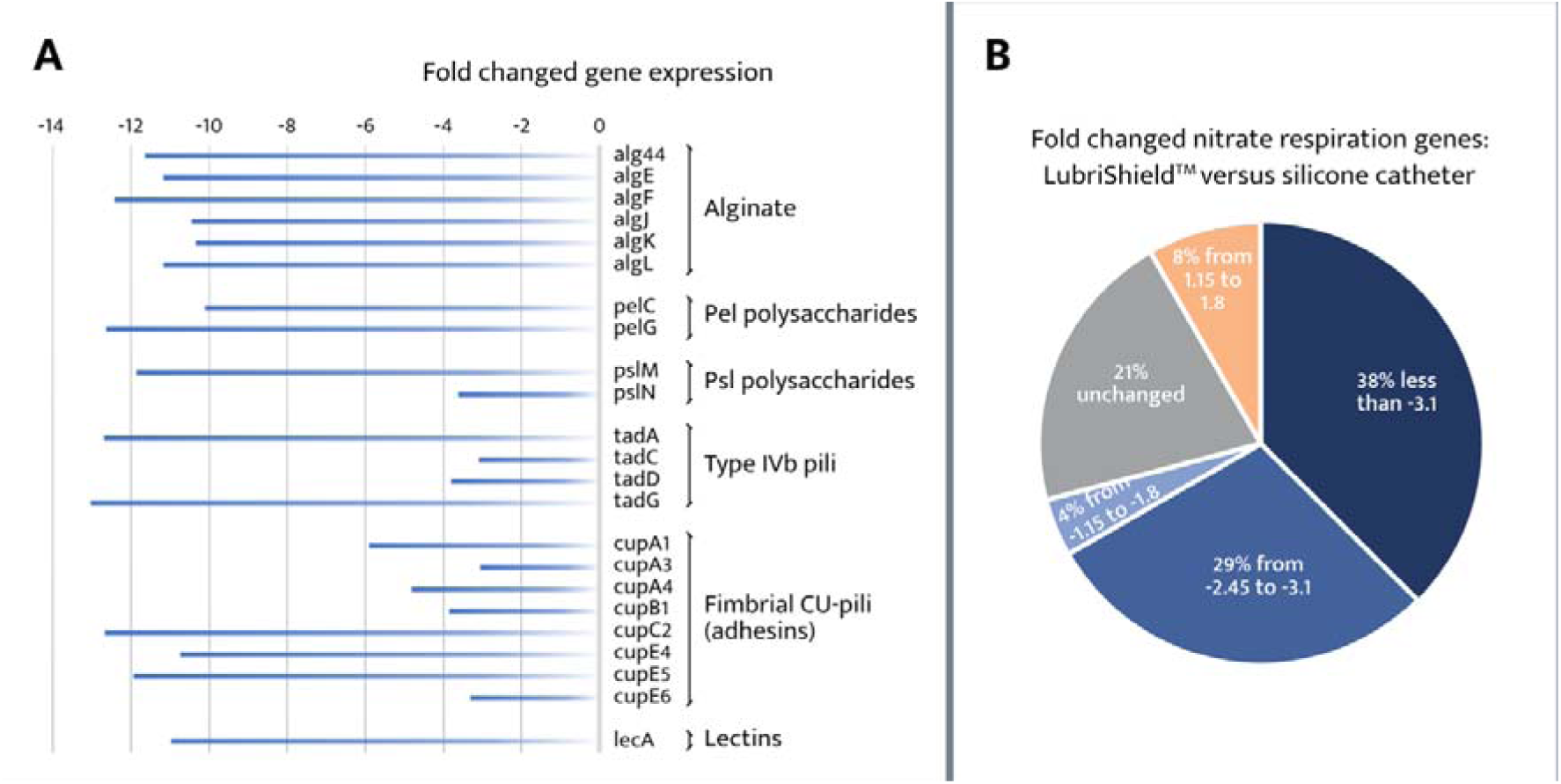
Transcriptome of surface-adhering bacteria after 7-day culture in AUM. **A.** Gene expression analysis of LubriShield^TM^ adhering P. aeruginosa showed significant down-regulation of biofilm-associated EPS operons compared to silicone. **B.** Genes associated with anaerobic nitrate respiration, a characteristic of biofilms, were significantly less expressed in LubriShield^TM^ adhering P. aeruginosa compared to silicone.

### Increased antibiotic response of bacteria present on LubriShield^TM^ coated catheters

To assess if the biofilm-preventive effect of the LubriShield^TM^ coating also included reducing antibiotic recalcitrance, two common biofilm-forming uropathogenic strains, one Gram-negative and one Gram-positive, were tested in long-term cultures in artificial urine on LubriShield^TM^ coated catheters compared to uncoated control catheters. Two different strategies were used: i) to treat with a bactericidal antibiotic (colistin) and then remove bacteria for subsequent viable count determination, ii) to directly assess metabolic status of the cells after a bacteriostatic antibiotic treatment (vancomycin), without removing cells from the catheter surface.

In the first case, after seven days with fresh daily media change at biofilm formation conditions, adhering *K. pneumoniae* were challenged with colistin (polymyxin E, 16 µg/mL) or left untreated for 2 hours and then assessed for viable counts. Results depicted in the boxplot figure 4A show a significantly improved response to treatment of bacteria present on the LubriShield^TM^ coated catheter compared to the uncoated catheter (fig. 4A, reduction from log_10_(3.2) to log_10_(4.7), n=3, Student’s t test).

**Figure 4.**
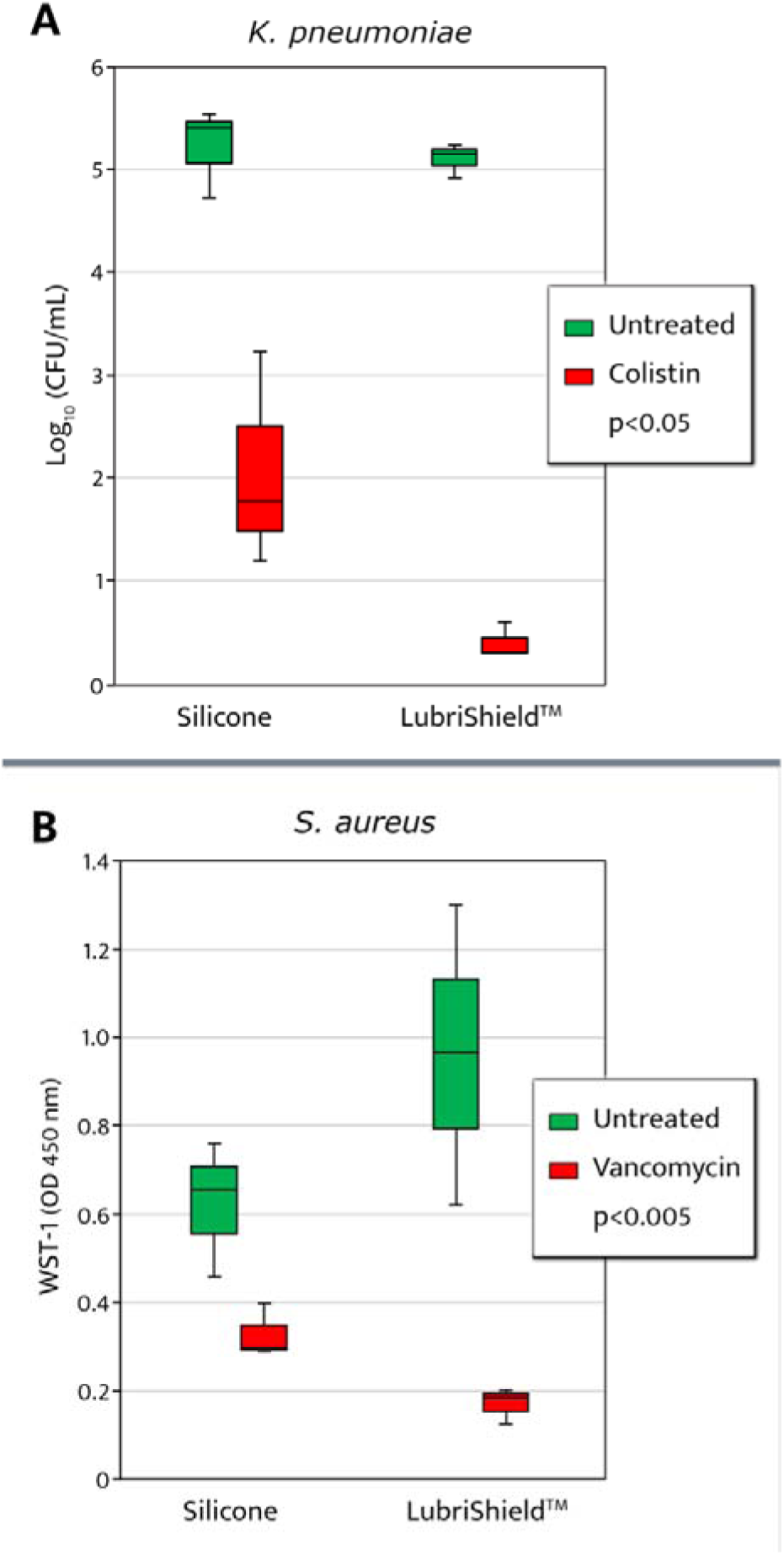
Treatment response to antibiotics after 7 days biofilm culture on catheters. A. Quantification of catheter-adhering K. pneumoniae by log_10_ (CFU/mL) count on uncoated Control versus LubriShield^TM^ coated catheters treated with colistin (red) or no treatment (green) (n=3). B. Evaluation of metabolic activity by measuring WST-1 of catheter-adhering S. aureus on uncoated control-versus LubriShield^TM^ coated catheters treated (green) with vancomycin or no treatment (red). P-values from Student’s t test comparing treatment-induced effect (n=3).

To test if the LubriShield^TM^ coating also resulted in an increase in the antibiotic responsiveness in Gram-positive bacteria, coated and uncoated control catheters were inoculated with *S. aureus*. After 7 days of culture in AUM, loosely bound bacteria were rinsed off and catheter pieces were treated for 2 hours with vancomycin (25 µg/mL) with identical untreated pieces serving as a reference. Reduction of the nontoxic tetrazolium salt, WST-1, which correlates to the cellular respiration rate, was then analyzed with a spectrophotometer ^23^. Results depicted in figure 4B show 33% lower WST-1 signal from the adhering bacteria on uncoated compared to LubriShield^TM^ coated catheter pieces (green boxes). This phenomenon is often associated with biofilm-living bacteria known to reduce their metabolism. After antibiotic treatment, 82% of the metabolic activity of the LubriShield^TM^ coated catheter-adhering bacteria was abolished, while a significantly less reduction (46%) was seen in bacteria present on the uncoated control-catheters (p<0.005).

### LubriShield^TM^ generated a permanent, non-release and nontoxic surface with low friction and high wettability

The instant the LubriShield^TM^ coated Foley catheter was wetted in either water or artificial urine medium it became highly lubricious. Assessment of the sliding friction of the coated catheter, revealed that the coefficient of friction (CoF) of *LubriShield*^TM^ coated samples was 28 times lower compared to the uncoated silicone catheter samples (CoF = 0.02 vs. 0.64 shown in fig. 5A, p<10^−7^).

**Figure 5.**
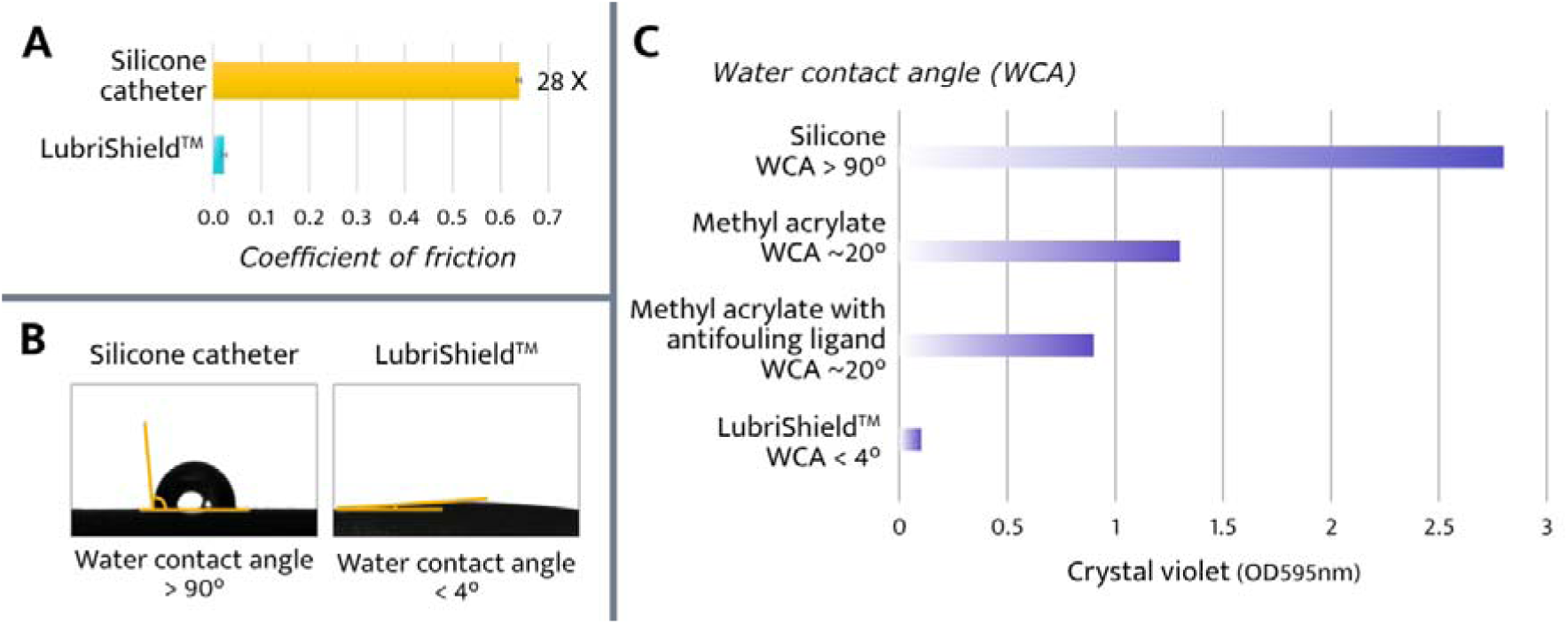
Physical characteristics of the coating. **A.** Sliding friction analysis of the catheter surface in a wetted state comparing uncoated standard silicone surface to the LubriShield^TM^coating is shown as the coefficient of friction. **B.** The LubriShield^TM^ coated catheter surface was tested for hydrophilicity and the water contact angle after wetting was measured and compared to uncoated silicone catheter surface.**C.** Biofilm assessment by crystal violet quantification of a 7-day culture of K. pneumoniae in artificial urine comparing uncoated silicone- and LubriShield^TM^ coated catheter surfaces to modified LubriShield^TM^ coatings with coupled hydrophobic methyl acrylate with or without our patented antifouling ligand.

We also assessed the wettability of the LubriShield^TM^ coating by the sessile drop technique, which measures the angle of which a water drop expands on a surface. This is called the water contact angle (WCA) and reflects the hydrophobicity of a material surface, where WCA larger than 90° is considered hydrophobic and smaller than 90° hydrophilic ^24^. When the water droplets spread promptly to completely wet the exposed area and make a compact water layer the surface is called superhydrophilic and has a WCA below 10°. The results from the sessile-drop technique performed by an external certified laboratory, Optas Ltd, is depicted in figure 5B and revealed that the LubriShield^TM^ coated catheter surface has a WCA of nearly 0°.

To demonstrate the critical role of the superhydrophilic nature of the LubriShield^TM^ coating and the incorporation of a proprietary antifouling ligand in its biofilm-preventive effect, two modified LubriShield^TM^ coated catheter samples were prepared. These samples included a hydrophobic monomer, methyl acrylate, either with or without the antifouling ligand. The modification increased the WCA of the sample surfaces to 20°. Biofilm formation by *K. pneumoniae* was quantified after 7 days culture in artificial urine to evaluate the impact of these modifications. As shown in figure 5C, the uncoated control silicone catheter with a hydrophobic surface (WCA of 100°) exhibited the highest level of biofilm formation, as measured by crystal violet staining. The methyl acrylate-modified coating (WCA∼20°) showed a 54% reduction in biofilm formation compared to the uncoated control. When the antifouling ligand was incorporated into the methyl acrylate-modified coating, biofilm formation was further reduced by an additional 30%. Notably, the unmodified LubriShield^TM^ coated catheter samples, with a superhydrophilic surface (WCA 0°), exhibited almost no biofilm formation under the same conditions (fig. 5C).

These findings highlight that the unique combination of the covalently bonded superhydrophilic polyacrylic acid hydrogel and the antifouling ligand produced a robust and durable antifouling effect, effectively preventing biofilm formation throughout the duration of use.

### LubriShield^TM^ is safe, does not release any toxic compounds and has no antimicrobial or cytotoxic effect

To assess the safety and biocompatibility of the LubriShield^TM^ coated catheter, we performed various *in vitro* tests. The chemical characterization of the coated catheter revealed that no chemical-substance release from the coating could be detected (ISO 10993 Part-18: 2020 (EJIAMD-1:2022 and ISO 10993-12:2021). In addition, long-term bacterial cultures on the LubriShield^TM^ coated catheter, showed no antimicrobial or bactericidal effect (fig. 1e) and thus did not induce any host reactions against toxic bacterial debris. Furthermore, cytotoxicity tests of the LubriShield^TM^ coating were performed on endothelial cells according to ISO standard 10993-5 and showed negligible cytotoxicity against human bladder and uroepithelial cells (table 1). The skin irritation assessment of the LubriShield^TM^ coating was also negative according to the ISO standard 10993-5 (table 1). These results confirmed that the LubriShield^TM^ coated Foley catheter was without toxicity, and compatible with biological systems.

**Table 1.** Cytotoxicity test result. Cytotoxicity testing according to ISO 10993-5

|  |  |
| --- | --- |
| *Cytotoxicity test of extract* | No cytotoxic potential |
| *EpiDerm skin irritation test* | No skin irritation potential |
| *Cytotoxic test of direct contact to L929 mouse cells* | No cytotoxic potential |
| *Cytotoxic test of direct contact to human uroepithelial/bladder cells* | No cytotoxic potential |

## Discussion

Catheter-associated urinary tract infections (CAUTIs) continue to be among the most prevalent healthcare-associated infections in both hospital and community care settings^25^. Even with the most scrupulous nursing care and rigorous application of sterile closed drainage systems, bacteriuria is almost inevitable in patients undergoing long-term catheterization (> 28 days), and this group of patients are particularly vulnerable to resulting complications ^26^. Currently, coatings on urinary indwelling catheters that release antimicrobial compounds (silver alloy or antimicrobials) have only been reported to yield positive results for short-term use.

Adherence of bacteria and fungi is a key step in infection; once adherent, the microorganisms can colonise and form biofilm often leading to CAUTI or blockage of the catheter lumen. It is therefore an urgent need to develop coatings on indwelling catheters that can prevent or reduce the formation of biofilms including crystalline biofilms. The common property of biofilms of different species is the presence of a matrix surrounding the community, shaping and protecting it, as well as, immobilizing its members. Microbial adhesion leading to biofilm formation is a complex process controlled by the interplay between physicochemical, mechanical, topographical surface properties, characteristics of the microorganisms, and environmental conditions. Most biofilm studies overlook the influence of surface parameters and surrounding hydrodynamic conditions on bacterial sensing and binding behaviour to a substrate. When bacteria adhere to a surface, the adhesion forces exerted by the substratum can induce nanoscopic deformation of the bacterial cell wall, leading to changes in the intracellular cytoplasmic pressure ^27,28^. Membrane-located sensor molecules subsequently react, which translates into a biological response with the synthesis and secretion of EPS ^27,28^. Superhydrophilic surfaces create a dense layer of water molecules, which weakens the interaction between cell surfaces and substratum material, thereby reducing cell adhesion ^29,30^. Studies have demonstrated that both Gram-positive and Gram-negative bacteria produce less EPS and are more susceptible to antibiotics on hydrophilic surfaces compared to hydrophobic surfaces with stronger adhesion properties ^31–33^. Similar behaviour has been observed with *C. albicans* biofilm formation on surfaces with high wettability ^34,35^. In addition to surface properties, hydrodynamic conditions, such as urine flow, play a critical role in biofilm development. The shear stress exerted by urine flow stimulates uropathogens to enhance their adhesion by producing thicker, more resistant biofilms as a countermeasure ^36^. Catheter encrustation caused by urease-producing organisms is also aggravated under these conditions and represents a serious complication for patients with indwelling catheters ^37^. As a result, indwelling urinary catheters provide an ideal environment for biofilm formation, significantly increasing the risk of recalcitrant clinical infections. In this study, we demonstrate the effectiveness of the LubriShield™ coating in preventing biofilm formation under both static and dynamic flow conditions.

Biofilm-related infections are notoriously resistant to a broad range of antibiotics and biocides, posing a significant clinical challenge due to persistent symptoms despite prolonged antimicrobial treatments. This recalcitrance is attributed to biofilm-specific features, including i) EPS-induced tolerance, ii) gene-regulated resistance mechanisms, and iii) dormant persister cells ^38–40^. In this study, we evaluated the susceptibility of catheter surface-associated bacteria after 7 days of culture in AUM to two common antibiotics used for treating complicated urinary tract infections. Our results demonstrated that biofilm-forming *K. pneumoniae* on LubriShield^TM^ coated surfaces were significantly more responsive to colistin, an antibiotic commonly used for multidrug-resistant Gram-negative infections, compared to uncoated silicone catheter surfaces. Similar results were shown for vancomycin, the therapy of choice for difficult-to-treat staphylococcal infections, which was significantly better at reducing the metabolic state in *S. aureus* after 7 days of culture on LubriShield^TM^ than on silicone catheter surface. These findings suggest that LubriShield^TM^ associated bacteria remain in a state comparable to planktonic cells, making them more susceptible to antibiotic treatment.

With growing evidence that biofilm formation plays a major role in medical device-related infections, updating detection standards to reflect advances in our understanding of biofilm composition is essential for accurate assessment and effective treatment. Existing standards, such as live/dead fluorescent staining, bioluminescent assays detecting ATP consumption, and culture-based analysis of colony-forming units (CFUs), primarily target metabolically active bacteria. These methods are therefore better suited for evaluating bactericidal antimicrobials. However, these assays often produce false-negative results when analyzing bacterial biofilms due to the presence of dormant persister cells with exceptionally low growth rates ^41^. To reliably differentiate planktonic bacteria from biofilm-forming bacteria, assays must target the extracellular polymeric substance (EPS) components, as planktonic bacteria do not produce EPS. The most widely used method is the colorimetric crystal violet (CV) assay, which binds to negatively charged components of the EPS – such as polysaccharides, proteins, extracellular DNA (eDNA), and lipids – as well as to live and dead bacteria. The CV assay’s simplicity, broad detection spectrum, and quantifiability make it ideal for *in vitro* biomass assessment. However, *in vivo* applications can be confounded by host-induced protein responses that interfere with results ^42^. Microscopic imaging techniques, such as episcopic differential interference contrast (EDIC) microscopy, provide an excellent way to examine the morphology of colonized surfaces *ex situ*, revealing the spread and structure of biomass sheaths or patches ^18^. Fluorescent tagging of specific EPS components further enhances biofilm detection, enabling precise confirmation of biofilm presence and composition. Finally, since biofilm formation and its emergent properties are regulated by genes responsive to surface sensing, stress, and quorum sensing signals, RNA-seq analysis is a powerful tool. It provides a detailed snapshot of the gene expression profile of the colonizing microbial population, offering critical insights into the biofilm-forming state ^7,43^.

The LubriShield^TM^ coating is designed to create a biocompatible, non-releasing, superhydrophilic, and permanent surface without exhibiting any microbial killing effects. Microbial incubation in artificial urine demonstrated significant inhibition of biofilm formation by the 8 most prevalent uropathogenic microbes under both static and dynamic conditions. The coating’s near-zero water contact angle was directly associated with its unique antifouling properties. Our findings suggest that interfering with bacterial surface-sensing mechanisms is critical for preventing the transition to a biofilm-forming state and the activation of biofilm-specific RNA expression profiles. Notably, no antimicrobial or bactericidal effects were observed in long-term cultures, and chemical characterization confirmed the absence of any substance release from the coating. In addition to its antifouling properties, the LubriShield™ coating exhibited a 28-fold reduction in surface friction compared to uncoated silicone catheters, further highlighting its potential to improve clinical outcomes by reducing mechanical irritation and biofilm formation.

Overall, these experimental findings suggest that the novel LubriShield^TM^ coating has the potential to significantly improve clinical outcomes by reducing patient discomfort, the risk of urethral stenosis, encrustation, and CAUTIs. Furthermore, the results indicate that catheter-related bacterial infections could potentially be treated with antibiotics without the need to remove the catheter. While these preclinical findings are promising, clinical studies are necessary to confirm their applicability in vivo. Ongoing Phase I and Phase II trials aim to evaluate the safety, efficacy, and clinical utility of LubriShield™ coated catheters in patient populations.

## Materials and methods

### LubriShield^TM^ coating

#### Medical devices

The coating was applied on a CE-marked medical grade silicone Foley catheter certified according to ISO 13485 (Sterimed Group). The uncoated silicone Foley catheter was used for all comparisons.

#### UV-induced polymerization

The grafting of silicone catheters was initiated via a UV-induced polymerization process. Prior to grafting, the catheters were dipped in a benzophenone (photo-initiator) ethanol solution. After drying, the catheters were immersed in an aqueous grafting solution containing acrylic acid under UV irradiation. Washing in deionized water and then ethanol was performed after the grafting process. The uniformity of the grafted surfaces of the silicone catheters was analysed by staining in an aqueous solution containing methanol crystal violet (4%). After thorough washing in deionized water the colouring of the coated silicone catheters was compared with a stained ungrafted silicone catheter (supplementary fig. S2).

#### Coupling of the proprietary antifouling ligand

The anti-fouling ligand, “2-(pyridyldithio)ethylamine (PDEA)”, was attached to the carboxylated surface by chemical coupling reactions essentially as reported in the literature ^44^. In short, the carboxylic groups in the grafted layer were activated with an aqueous solution of 1-ethyl-3-(3-dimethylaminopropyl) carbodiimide (EDC) and *N*-hydroxysuccinimide (NHS) and then reacted with PDEA. Intermittent washing in deionized water was performed for each reaction. Chemical characterization of the coated catheter revealed that no chemical-substance release from the coating could be detected (ISO 10993 Part-18: 2020 (EJIAMD-1:2022 and ISO 10993-12:2021).

### Microbiology

#### Materials

The following materials were used for the microbiological study work : Luria broth and Luria agar plates (Karolinska Hospital Substrate unit, Sweden), phosphate buffered saline (PBS, Karolinska Hospital Substrate unit, Sweden), crystal violet (CAS 548-62-9, Acros Organics, USA), EbbaBiolight 680 (Ebba Biotech, Sweden), concanavalin A (Alexa Fluor® 594 conjugate, Thermo Fischer Scientific, USA), FilmTracer™ SYPRO™ Ruby Biofilm Matrix Stain (Thermo Fischer Scientific, USA), colistin sulphate (CAS 1264-72-8, MedChemExpress, USA), vancomycin hydrochloride (CAS 1404-93-9, MedChemExpress, USA). Artificial urine medium (AUM) was freshly prepared according to Brooks T. et al. 1997 every day in the lab, pH adjusted to 6.5 and filter sterilized before use ^45^. Chemicals used for AUM preparation are listed in supplementary table S2.

#### Strains used

A set of microorganisms, representing Gram-positive and Gram-negative bacteria species, as well as *Candida albicans*, was used (supplementary table S3).

#### Culturable count

Culturable count was done by counting microorganism colony forming units. For bacteria in liquid medium, a serial 10x dilution in 1 mL of PBS was done. 100 µL of resulting microorganism suspensions was placed on Luria agar plates, spread with sterile glass beads and incubated at 37⁰C overnight. Colonies were counted and CFU/mL in original culture calculated. For surface-attached bacteria, catheter pieces were placed in 1.5 mL of PBS in a 2-mL microcentrifuge tube and sonicated with Cell Disruptor (AOSENR, China) 30 seconds sonication/ 10 minutes cooling / 30 seconds sonication to remove attached bacteria and disrupt biofilm. Resulting suspension was used for serial dilutions and colony counting as described above.

#### Static biofilm growth test

The test was performed on 2-cm catheter pieces placed into 2 mL-microcentrifuge tubes with only shaking creating shear force necessary for the biofilm formation. 1 ml of freshly made AUM containing resuspended 10^9^ CFU/mL for bacteria or 10^8^ CFU/mL for *Candida* was added. Tubes were incubated on a rotary shaker (90 rpm) at 37⁰C and media was changed daily. After seven days or more, the pieces were removed, gently rinsed and analysed.

#### Dynamic biofilm growth test with flow model

The test was performed on 7-cm catheter sections placed in line with a peristaltic pump for a continuous medium flow. *P. mirabilis,* 10^9^ CFU/mL, in freshly made AUM was seeded inside the catheters for 30 minutes before connecting to the system and initiating flow. Cultures were kept at 37°C and continuously supplied with fresh medium at a rate of 1 mL/minute. After 3 days, the sections were removed, gently rinsed and analysed.

#### Dynamic biofilm growth test with glass bladder model

The model consists of a glass vessel maintained at 37°C by a water jacket (fig. 2B, left panel). The model was sterilized by autoclaving and then a catheter was inserted aseptically into the vessel through a section of silicone tubing attached to a glass outlet at the base. The catheter balloon was inflated with 10 mL of water, securing the catheter in position and sealing the outlet from the bladder. The catheter was then attached to a drainage tube and a reservoir bag located below the level of the glass vessel. Sterile AUM was supplied to the bladder via a peristaltic pump. In this way a residual volume of urine collected in the bladder below the level of the catheter eyehole. As AUM was supplied to the model, the overflow drains through the catheter to the collecting bag. Six models with three LubriShield^TM^ coated catheters and three uncoated catheters were set up to study the biofilm formation of *P. aeruginosa*, responsible for about 10% of all CAUTI ^46^. An inoculum of 100 µL of 10^9^ CFU/mL of *P. aeruginosa* in AUM was added to the residual volume in the vessel for 1 h to establish itself before the supply of AUM, at a rate of 1.5 mL/min, was switched on. After five days the supply of AUM to the bladder was turned off, the balloon was deflated, and the catheter was removed through the base of the model.

#### Crystal violet staining

Crystal violet staining was done as described in the literature ^47^. Adaptation from Kanematsu H and Barry DM. Springer Nature. 2020 (Chapter 6.1, page 114) and Colomer-Winter C et al. Bio-protocol. 2019 (page 8) with modifications ^48,49^. Briefly, catheter samples were immersed in 1 mL of 0.04 % crystal violet solution in PBS for 10 min, followed by three rinses in PBS with blotting towards paper in between and left to dry. For visualization, the catheter surface was pressed against Scotch® mending tape (3M Japan, Tokyo, Japan) to transfer the biofilm. Structure of the transferred biofilm on the flat surface of the Scotch® mending tape was visualized directly or under the microscope attached to the sample glass. For quantification, biofilm was transferred on a wet (PBS) cotton swab by gently scratching the surface of the catheter piece. The head of the cotton swab was then removed and placed into 1.5 mL of 96% ethanol solution for crystal violet extraction (2h, room temperature) before spectrophotometric quantification at OD595 nm.

#### EDIC microscopy

Qualitative assessment of biofilm development. The catheter sections were examined using an episcopic differential interference contrast (EDIC)/epifluorescence (EF) Nikon Eclipse LV100D microscope (Best Scientific, UK) using a metal halide light source (EXFO X-CITE 120 fluorescence system), long working distance metallurgical objectives (Nikon Plan Achromat) and a high-resolution camera (QImaging Retiga EXi Cooled Digital CCD monochrome camera with RGB colour filter module). Images were captured and processed using ImagePro image capture software. Experiments were repeated three times with two sections (from each experiment) used for EDIC microscopy at each time point. The entire length of the sections was examined using a x 50 objective with representative images being taken over 10–50 fields of view (including at the higher magnification of × 1000) ^18^.

#### Fluorescence microscopy

For catheter surface analysis, 1 mL staining solutions of either concanavalin A (10 μg/mL, Alexa Fluor® 594 conjugate) or EbbaBiolight 680 (0.1 μg/mL) were prepared in PBS. FilmTracer™ SYPRO™ Ruby Biofilm Matrix Stain was purchased ready to use. Catheter pieces were stained for 30 min in the dark and then washed twice for 5 minutes in PBS with paper blotting in between. Fluorescent microscopy was conducted using a Trinocular Compound EPI-Fluorescence Microscope Model M837FLR (OMAX, USA) in the red and green channel mode.

#### RNA-seq analysis

For the RNA-seq analysis, *Pseudomonas aeruginosa* PA01 strain was grown on a surface of 2-cm catheter pieces (16Fr, LubriShield^TM^ vs uncoated silicone catheter) in 1 mL of AUM in microcentrifuge tubes for 7 days at 37⁰C with slow shaking and daily changing of the medium. Catheter pieces were gently removed and thoroughly rinsed 5 times in PBS with liquid carefully removed in between. After that, they were frozen in pre-chilled 2-cm tubes and stored at −80⁰C or in dry ice during transportation prior to the RNA extraction procedures. RNA extraction, library construction, prokaryotic transcriptome sequencing was performed by BGI Genomics (Hong Kong, China). Total RNA was isolated. The library was constructed in the following steps: a) rRNA depletion, b) RNA fragmentation, c) first cDNA strand synthesis, d) second cDNA strand synthesis, e) end repair and adaptor ligation, f) PCR amplification, g) library purification and qualification. After that sequencing was performed using DNBSEQ sequencing platform. Sequencing reads were filtered, mapped to the published Pseudomonas aeruginosa PA01 genome software ^50^ using HISAT2 ^51^ and StringTie software for assembly ^52^. Bowtie2 was used to align clean reads to the reference sequence ^53^, and RSEM was used to calculate the expression of genes and transcripts ^54^. For the differential analysis between groups, DEGseq method was used ^55^ and between samples Poisson distribution method was used ^56^. Hierarchical clustering analysis of DEGs, gene ontology analysis of DEGs and pathway analysis of DEGs was performed. For mRNA, annotation following databases were used: NR, NT, GO, KOG, KEGG, Uniprot, COG, PRG, String, TF. For lncRNA, the following annotated databases were used: miRBase, Rfam. Hierarchical clustering analysis of DEGs, gene ontology analysis of DEGs and pathway analysis of DEGs was performed.

#### Antibiotic sensitivity assays

For the antibiotic sensitivity assay 7-day biofilm cultures of bacteria were prepared on LubriShield^TM^ coated vs. uncoated 2-cm catheter pieces as it is described above. The pieces were thoroughly washed from loosely adhered bacteria (3x, 1 min washes in 1 mL of PBS with paper blotting in between and placed into 1 mL of fresh AUM to recover for 1h at 37⁰C with shaking (90 rpm). a) For the colistin sensitivity assay: three pieces of each kind were then treated with colistin (16 μg/mL in AUM, 37⁰C, 2h), while another triplicate was used as the untreated reference. After the treatment, Cell Disruptor (AOSENR, China) was used to remove adhered bacteria and disrupt biofilm (30 seconds sonication / 10 minutes cooling / 30 seconds sonication). 10x serial dilutions were made and 100 µL of each of them were placed on Luria agar plates and spread with sterilized glass beads. Next day, colonies were counted and CFU/mL in the original suspension calculated. b) For the vancomycin sensitivity assay: three pieces of each kind were treated with vancomycin (25 μg/mL in AUM, 37⁰C, 2h), while another triplicate was used as the untreated reference. 50 µL of the WST working solution was added to each of the tubes and incubated for the additional 1h at 37⁰C. 400 µL sample was then spun down to remove debris and OD450 nm measured with spectrophotometer.

#### Water Contact Angle visualization and measurement

Measurement of the water contact angle, WCA, by sessile drop imaging analysis was performed by Optas LTD (UK), an independent expert house. The samples were pre-wetted in HPLC Grade water before measuring the contact angle at the water – air – film surface interface with a tracking oscillating drop tensiometer (Teclis Scientific).

#### Sliding friction measurement

Measurements of the dynamic coefficient of friction was performed by Optas LTD (UK). The samples were pre-wetted in HPLC Grade water before a sled weight of 90 g was applied. After a delay of three seconds a test speed of 24 mm /minute was applied, for a distance of 20 mm. The measurements were performed according to ISO 8295.

#### Statistical analysis

Biofilm formation measured by crystal violet comparing LubriShield^TM^ catheter surface with control silicone catheter surface was analysed by Student’s unpaired, two-tailed t-test. Viability data by CFU/mL were log-transformed to achieve a normal distribution. The log reduction of each experiment was determined by subtracting the log CFUs before treatment from the log CFUs after treatment. Data are plotted as the means with staple bars showing the standard error of the mean (SEM). The probability value less than 0.05 was considered significant.

## Acknowledgement

We extend our gratitude to the CytaCoat AB team for their valuable contributions and technical expertise. In particular, we thank Å. Wang and N. Asadli for their dedicated work in coating and validating Foley catheters, and E. Roca for her meticulous handling of the microbial strains. This research was generously supported by CytaCoat AB.

