## Supplementary figure 1 for "*LubriShield^TM^* - a unique permanent coating for indwelling urinary catheters that impedes surface-associated uropathogens from forming biofilm"

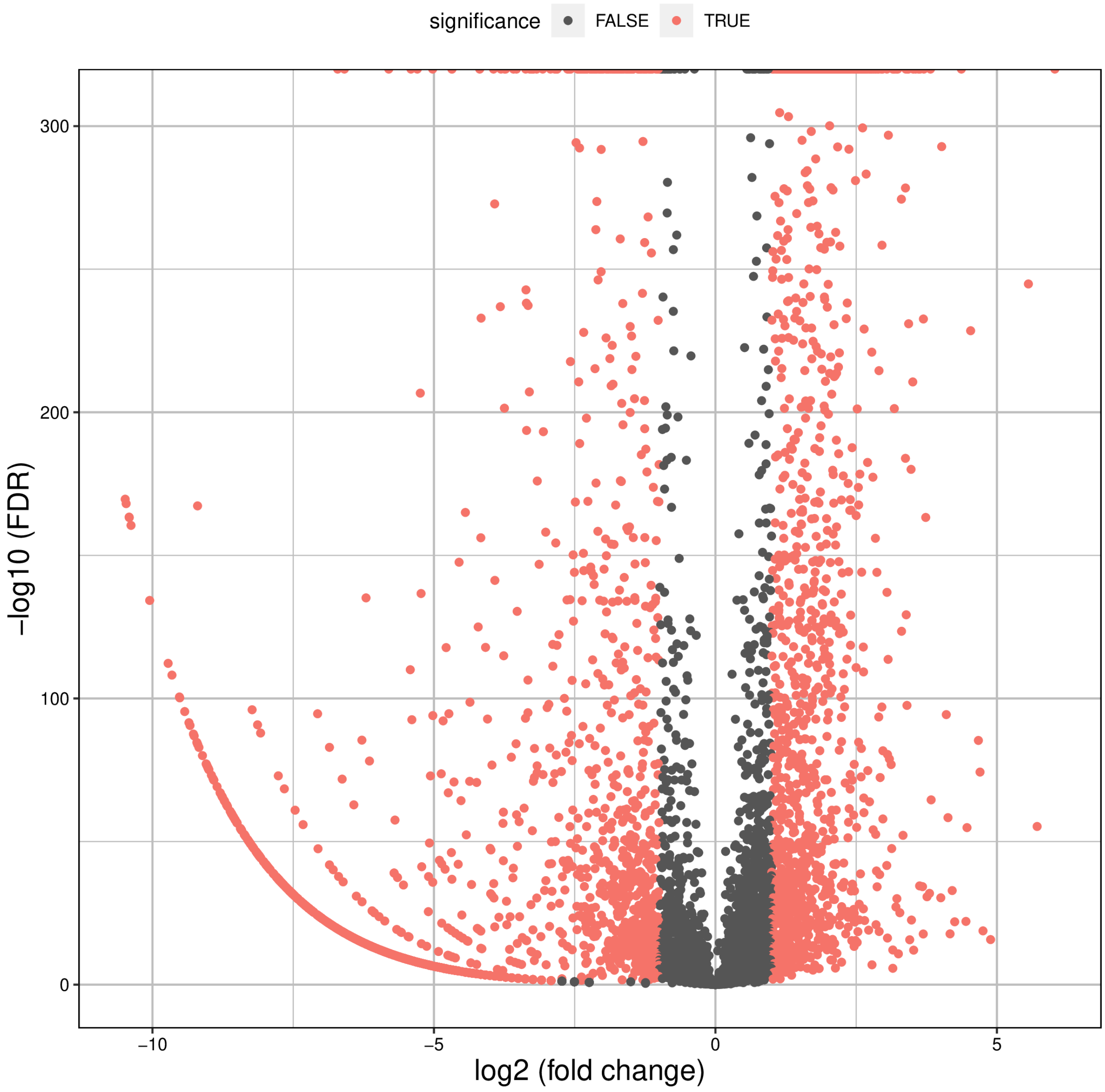


***Figure S1. Volcano plot representing all expressed transcripts.*** For every transcript, the fold change of LubriShield™ versus silicone catheter-associated *P. aeruginosa* was plotted against the -log P value. Statistically significant differentially expressed genes, with a fold change ≥1.5 or ≤ -1.5, are depicted as red, insignificant as black dots.
