## Supplementary table 1 for "*LubriShield^TM^* - a unique permanent coating for indwelling urinary catheters that impedes surface-associated uropathogens from forming biofilm"

**Table S1. Fold change in gene expression related to nitrate respiration, arginine fermentation and other hypoxia-related genes in LubriShield™ versus silicone catheter-associated P. aeruginosa**

| | Gene ID | Gene name | Gene description | Fold change |
|---|---|---|---|---|
| Nitrate respiration related genes | PA4922 | azu | azurin precursor | -1,9 |
| | PA2664 | fhp | flavohemoprotein | -4,68 |
| | PA3875 | narG | respiratory nitrate reductase alpha chain | -3,24 |
| | PA3874 | narH | respiratory nitrate reductase beta chain | -3,72 |
| | PA3872 | narI | respiratory nitrate reductase gamma chain | -4,19 |
| | PA3873 | narJ | respiratory nitrate reductase delta chain | -3,3 |
| | PA3877 | narK1 | nitrite extrusion protein 1 | -3,74 |
| | PA3876 | narK2 | nitrite extrusion protein 2 | -6,7 |
| | PA3879 | narL | two-component response regulator NarL | -2,34 |
| | PA3871 | nifM | probable peptidyl-prolyl cis-trans isomerase, PpiC-type | -5,29 |
| | PA0510 | nirE | uroporphyrinogen-III C-methyltransferase | -2,85 |
| | PA0516 | nirF | heme d1 biosynthesis protein NirF | -2,03 |
| | PA0512 | nirH | Siroheme decarboxylase NirH subunit | -1,94 |
| | PA0511 | nirJ | heme d1 biosynthesis protein NirJ | -1,91 |
| | PA0518 | nirM | cytochrome c-551 precursor | -2,31 |
| | PA0520 | nirQ | regulatory protein NirQ | -2,47 |
| | PA0519 | nirS | nitrite reductase precursor | -1,7 |
| | PA0524 | norB | nitric-oxide reductase subunit B | -2,34 |
| | PA0525 | norD | probable dinitrification protein NorD | -3,17 |
| Arginine fermentation related genes | PA5171 | arcA | arginine deiminase | -2,57 |
| | PA5172 | arcB | ornithine carbamoyltransferase, catabolic | -2,39 |
| | PA5173 | arcC | carbamate kinase | -3,07 |
| | PA5170 | arcD | arginine/ornithine antiporter | -2,41 |
| | PA0899 | aruB | N2-Succinylarginine dihydrolase | -2,57 |
| | PA0901 | aruE | N-Succinylglutamate desuccinylase | -1,01 |
| Other hypoxia-related genes | PA5427 | adhA | alcohol dehydrogenase | -2,37 |
| | PA2127 | cgrA | cupA gene regulator A, CgrA | -2,02 |
| | PA0459 | clpC | probable ClpA/B protease ATP binding subunit | -1,26 |
| | PA3126 | ibpA | heat-shock protein IbpA | -1,81 |
| | PA4236 | katA | catalase | -1,67 |
| | PA0835 | pta | phosphate acetyltransferase | -1,23 |
| | PA5495 | thrB | homoserine kinase | -2,92 |
| | PA0310 | | Fe2OG dioxygenase domain-containing protein | -2,08 |
