## Supplementary figure 2 for "*LubriShield^TM^* - a unique permanent coating for indwelling urinary catheters that impedes surface-associated uropathogens from forming biofilm"

Standard silicone catheter

LubriShield™ catheter

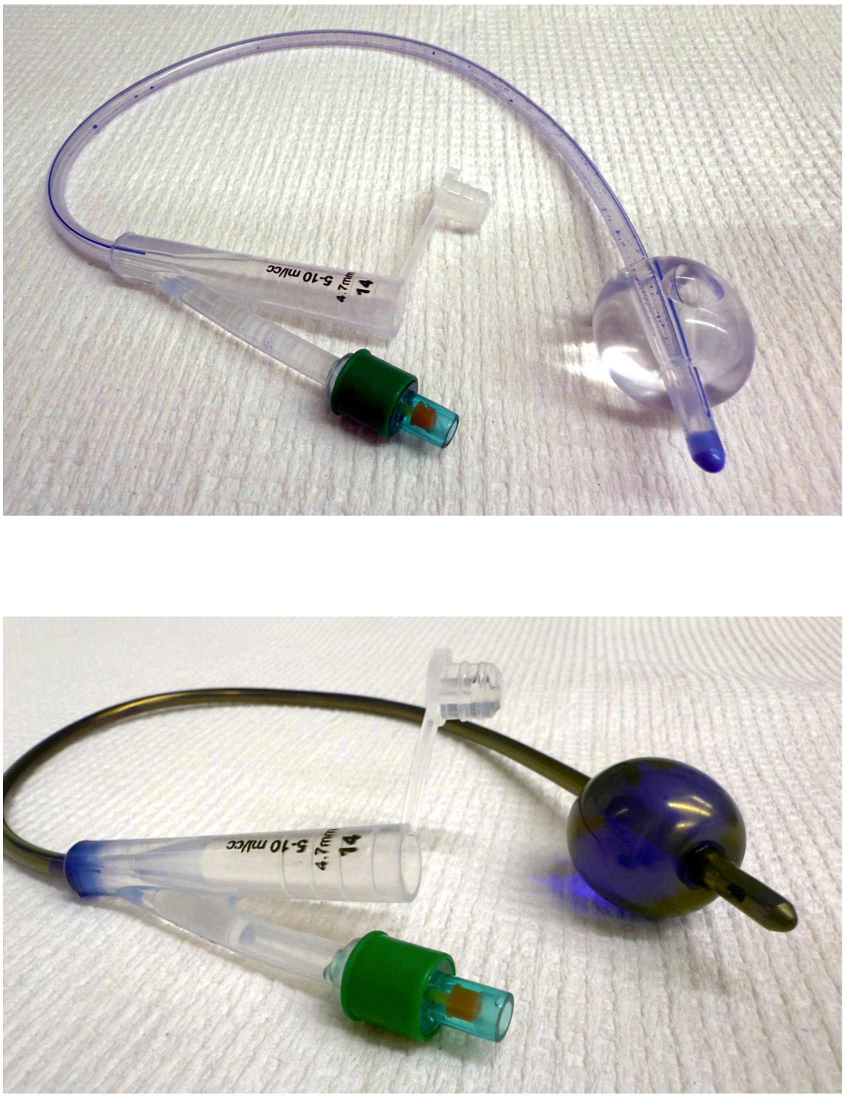

**Figure S2. Crystal violet staining of an uncoated and a coated Foley catheter.** The uniformity of the grafted surfaces of the silicone catheters was analysed by staining in an aqueous solution containing methanol and Crystal violet (4%).
