## Supplementary table 2 for "*LubriShield^TM^* - a unique permanent coating for indwelling urinary catheters that impedes surface-associated uropathogens from forming biofilm"

**Table S2. Chemicals used for artificial urine medium preparation**

| Chemical | CAS | Producer |
|---|---|---|
| Urea | 57-13-6 | Carlo Erba, Italy |
| Magnesium sulfate anhydrous | 7487-88-9 | Carlo Erba, Italy |
| Potassium phosphate monobasic | 7778-77-0 | Carlo Erba, Italy |
| Creatinine | 10028-24-7 | Ambeed, USA |
| Citric acid | 77-92-9 | neoFroxx, Germany |
| Yeast extract | 8013-01-02 | Thermo Scientific, USA |
| Calcium chloride dihydrate | 10035-04-8 | Thermo Scientific, USA |
| Sodium bicarbonate | 144-55-8 | Thermo Scientific, USA |
| Iron(II) sulfate heptahydrate | 7782-63-0 | Thermo Scientific, USA |
| Sodium chloride | 7647-14-5 | Thermo Scientific, USA |
| Ammonium chloride | 12125-02-9 | Thermo Scientific, USA |
| Potassium phosphate dibasic | 7758-11-04 | Thermo Scientific, USA |
| Sodium sulfate decahydrate | 7727-73-3 | Thermo Scientific, USA |
| Bacto™ Peptone | | Thermo Scientific, USA |
| DL-Lactic acid | 50-21-5 | TCI, Japan |
| Uric acid | 69-93-2 | TCI, Japan |
