## Supplementary table 3 for "*LubriShield^TM^* - a unique permanent coating for indwelling urinary catheters that impedes surface-associated uropathogens from forming biofilm"

**Table S3. Bacterial and fungal strains used in the study**

| Strain name | Description |
|---|---|
| *Pseudomonas aeruginosa PA01* | Wound isolate *P. aeruginosa*. Common laboratory strain |
| *Staphylococcus aureus B5381* | Clinical isolate *S. aureus* (Karolinska institute, Sweden) |
| *Klebsiella pneumoniae AO15200* | Uropathogenic *K. pneumoniae* strain |
| *Escherichia coli CFT073* | Uropathogenic *E. coli* strain |
| *Proteus mirabilis CCUG33828* | Uropathogenic *P. mirabilis* strain |
| *Enterococcus faecium #11* | Clinical isolate *E. faecium* (Karolinska institute, Sweden) |
| *Enterococcus faecalis #26* | Clinical isolate *E. faecalis* (Karolinska institute, Sweden) |
| *Staphylococcus epidermidis Se19* | Isolated from human peritonitis |
| *Candida albicans CCUG70317* | Uropathogenic *C. albicans* strain |
